# The WIPI homolog Atg18 binds to and tethers membranes containing phosphatidylinositol-3,4,5-triphosphate

**DOI:** 10.1101/2025.01.31.635529

**Authors:** Fernando L. Noguera-Cañas, Aitor Díaz-Agustín, Jose A. Gavira, Jose M. Paredes, Rafael Salto, Angel Orte, Francisco Conejero-Lara, Mercedes Martín-Martínez, Ángel Pérez-Lara

**Affiliations:** Department of Physical Chemistry, University of Granada, Campus Universitario de Cartuja, 18071 Granada, Spain; Instituto de Química Médica (IQM-CSIC), Juan de la Cierva 3, 28006 Madrid, Spain; Laboratory of Crystallographic Studies, (IACT-CSIC), Avenida de las Palmeras 4, 18100 Armilla, Spain; Department of Biochemistry and Molecular Biology II, University of Granada, Campus Universitario de Cartuja, 18071 Granada, Spain; Departamento de Química Física, Universidad de Granada. Avda Fuentenueva, 18071 Granada, Spain

**Keywords:** autophagy, PROPPIN, WIPI, membrane-protein interactions, polyphosphoinositides

## Abstract

Atg18 —for Autophagy-related gene 18— is a member of the PROPPIN (β-<u>prop</u>ellers that bind <u>p</u>olyphospho<u>in</u>ositides) family, which is known for its binding to phosphorylated phosphoinositides through two conserved binding sites. Although Atg18 binding to polyphosphoinositides is crucial for its roles in both autophagic and non-autophagic functions within cells, the precise molecular mechanism by which Atg18 selectively binds to specific phosphatidylinositols remains unresolved. Here, we combined molecular dynamic simulations and biophysical methods —including isothermal titration calorimetry (ITC), cosedimentation, fluorescence resonance energy transfer (FRET), and dynamic light scattering (DLS)— to characterize the interaction between Atg18 and polyphosphoinositides. In contrast to previous findings, we demonstrate that Atg18 binds to and clusters liposomes containing phosphatidylinositol-3,4,5-triphosphate (PtdIns(3,4,5)P_3_), suggesting that Atg18 oligomerizes and tethers opposing membranes containing this physiological phosphatidylinositol. Hence, our results provide new insights into how Atg18 and its mammalian homologs, WIPI —for WD-repeat domain phosphoinositide-interacting— proteins, may regulate organelle membrane organization and vesicle trafficking required for both autophagic and non-autophagic functions.

## Introduction

Atg18 belongs to the protein family of β-<u>prop</u>ellers that bind <u>p</u>olyphospho<u>in</u>ositides (PROPPIN), a family that plays crucial roles in the regulation of membrane dynamics and vesicular trafficking in yeast (1). Its primary structure consists of multiple WD40 repeats that assemble into a 7-bladed β-propeller fold, which contains a characteristic FRRG motif as part of two conserved binding sites for physiological phosphatidylinositols (PtdInsP_x_) (2, 3). The current view of the field is that Atg18 preferentially binds to phosphatidylinositol-3-phosphate (PtdIns(3)P) for its autophagic functions, such as the expansion of the autophagosomal membrane, and to PtdIns(3,5)P_2_ for non-autophagic functions, such as regulating PtdIns(3,5)P_2_ levels at the vacuole, retrograde transport from the vacuole to the Golgi, and membrane fission (1, 3-6). Furthermore, it is widely accepted that binding site I binds with a higher affinity for PtdIns(3,5)P_2_, whereas binding site II exhibits a higher affinity for PtdIns(3)P (7).

Despite recent advances on the Atg18 structure (8) and its interaction with PtdInsP_x_ (4, 9-11), the molecular mechanism by which the two binding sites of Atg18 bind stereoselectively PtdIns(3)P and PtdIns(3,5)P_2_ remains unclear. To elucidate the polyphosphoinositide binding mechanism of Atg18 and to determine the individual contributions of each phosphate group on the inositol ring to the stereoselectivity of Atg18, we performed both *in silico* and *in vitro* experiments using different PtdInsP_x_. Strikingly, our results reveal that Atg18 binds to and clusters vesicles containing phosphatidylinositol-3,4,5-triphosphate (PtdIns(3,4,5)P_3_), contradicting earlier findings (3, 4, 11, 12) and suggesting new physiological roles for Atg18.

## Results and Discussion

To investigate the Atg18 stereoselectivity for PtdInsP_x_, a series of *in silico* simulations were carried out. Induced fit docking (IFD) studies showed that Atg18 binds to inositol-1,3,4,5-tetrakisphosphate (Ins(1,3,4,5)P_4_) —the head group of PtdIns(3,4,5)P_3_— at binding sites I and II, and subsequent 500 ns molecular dynamic (MD) simulations revealed a stable binding of Ins(1,3,4,5)P_4_ at site I (Fig. 1 *A* and *B*). The analysis of the interactions at binding site I showed salt bridges and hydrogen bonds with phosphate 5 and phosphate 3, as expected (Fig. 1*A*). Moreover, simulation also revealed hydrogen bonds between Arg222 side chain and the non-bridging oxygens of phosphate 4, suggesting that Atg18 binds to PtdIns(3,4,5)P_3_, in contrast to previous *in vitro* results reporting no binding (3-5, 11, 12).

**Figure 1.**
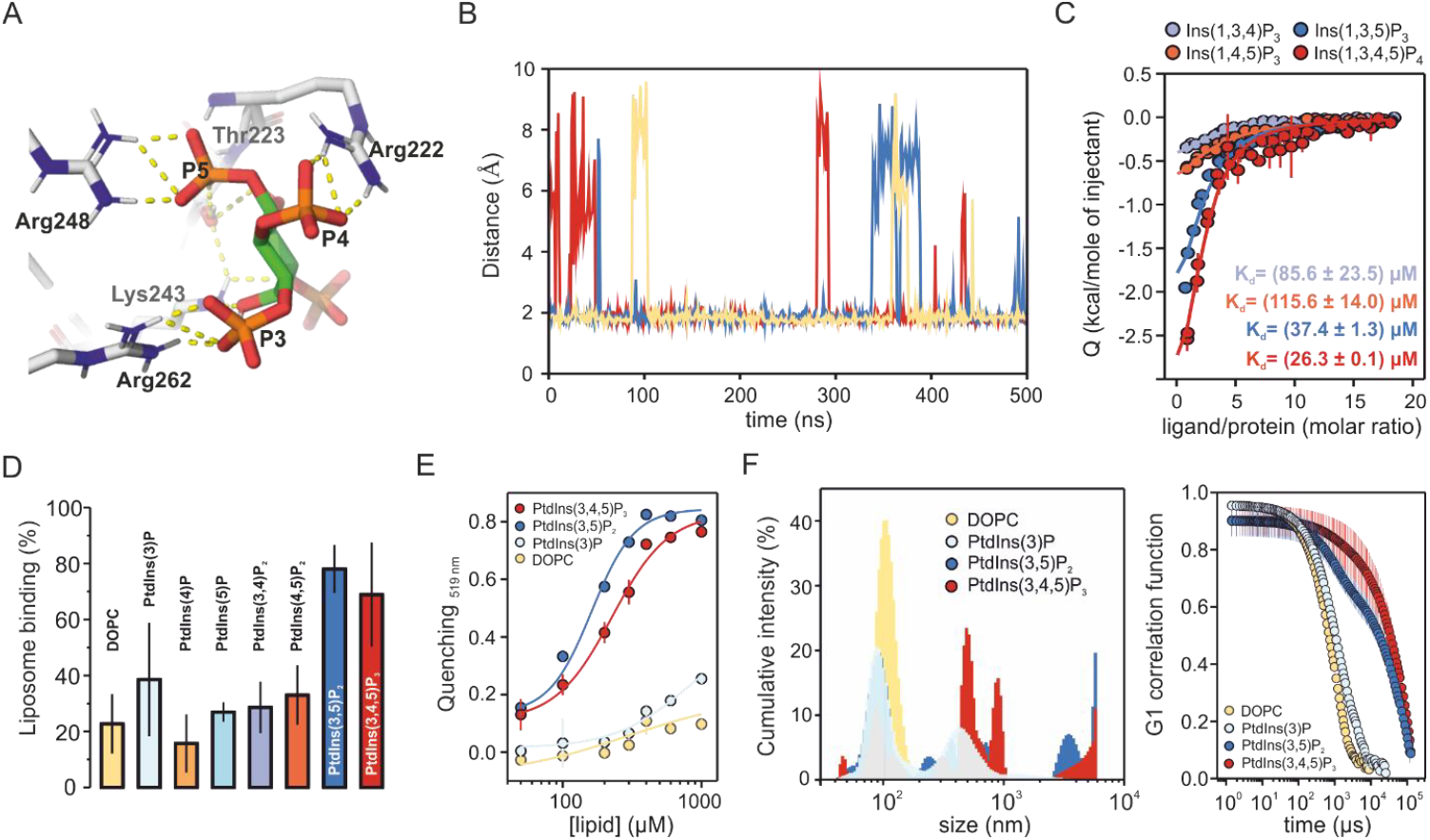
Atg18 binds to and clusters PtdIns(3,4,5)P_3_ containing membranes. (A) Representative 500 ns molecular dynamic conformation of Ins(1,3,4,5)P_4_ into the binding site I of Atg18 showing the salt bridges and hydrogen bonds (indicated by dashed lines) with phosphate 3 (P3), phosphate 4 (P4) and phosphate 5 (P5), while phosphate 1 does not show polar interactions. (B) Plot of the global distances between the NH_2_ protons of Atg18 Arg222 side chain and the non-bridging oxygens of Ins(1,3,4,5)P_4_ phosphate 4 over the time course of three 500 ns MD replicas (each represented by one colour) (C) Binding of Atg18 to phospholipid head groups of physiological polyphosphoinositides measured by ITC. 25 μM of Atg18 was titrated with main InsP_3-4_ stereoisomers at 25 °C in a buffer consisting of 50 mM HEPES and 150 mM KCl, pH 7.5. (D) Binding of Atg18 to large unilamellar vesicles (LUV) monitored by cosedimentation using sucrose-loaded liposomes containing 2% of physiological phosphatidylinositol (PInsP_x_) and 98% of 1,2-dioleoyl-sn-glycero-3-phosphocholine (DOPC). (E) Observed changes of the fluorescence (see *SI Appendix*) at 519 nm for the Alexa Fluor 488-labelled Atg18 after addition of liposomes containing the FRET acceptor Texas Red-DHPE. (F) Vesicle clustering by Atg18. DLS profiles and autocorrelation functions of the DLS measurements of LUVs (around 100 nm diameter) in the presence of 1 μM Atg18. All liposomes consisted of 2% of either PtdIns(3)P, PtdIns(3,5)P_2_, or PtdIns(3,4,5)P_3_, and 98% DOPC.

To further validate our *in silico* results, we performed isothermal titration calorimetry (ITC) experiments using previously studied PtdInsP_x_ head groups (InsP_x_). Our reasoning behind this approach was based on the idea that the head groups of PtdInsP_x_ enabled us to elucidate the stereoselectivity of Atg18 while minimizing potential interferences caused by other phospholipids and non-specific electrostatic or hydrophobic interactions between the protein and the membrane. Upon injecting inositol bisphosphates —Ins(1,3)P_2_, Ins(1,4)P_2_, and Ins(1,5)P_2_— into a solution containing Atg18, we observed only small changes in the binding enthalpy, thereby hindering quantitative assessment of the thermodynamic parameters of the binding process. In contrast, the binding of inositol tri- and tetrakis-phosphates —Ins(1,3,4)P_3_, Ins(1,3,5)P_3_, Ins(1,4,5)P_3_, and Ins(1,3,4,5)P_4_— to Atg18 was exothermic, with affinities ranging from tens to hundreds μM (Fig. 1*C*). However, because of their smaller size and lack of the larger hydrophobic tail, InsP_x_ have higher degrees of freedom and less steric hindrance than their corresponding PtdInsP_x_. Thus, to unambiguously confirm that Ins(1,3,4,5)P_4_-binding does also involve Atg18-PtdIns(3,4,5)P_3_ binding we performed a cosedimentation assay using liposomes containing different PtdInsP_x_. Our cosedimentation experiments confirmed that Atg18 binds to PtdIns(3)P and PtdIns(3,5)P_2_, showing higher binding for PtdIns(3,5)P_2_ (Fig. 1*D*). Interestingly, although previous studies reported that Atg18 binds exclusively to PtdIns(3)P and PtdIns(3,5)P_2_, and not to PtdIns(3,4,5)P_3_ (3, 4), our results revealed that Atg18 also binds to PtdIns(3,4,5)P_3_ with a binding comparable to that for PtdIns(3,5)P_2_ (Fig. 1*D*). To test these opposite results, we used a FRET assay (8) to determine the affinity of Atg18 to PtdIns(3)P, PtdIns(3,5)P_2_, and PtdIns(3,4,5)P_3_. Similar apparent dissociation constants (K_d_) were observed for liposomes containing PtdIns(3,5)P_2_ (K_d_ = 161 ± 17 μM) or PtdIns(3,4,5)P_3_ (K_d_ = 226 ± 26 μM), confirming our cosedimentation results (Fig. 1*E*). Additionally, Atg18 also bound liposomes containing PtdIns(3)P but with significantly lower affinity that it made difficult to accurately determine its K_d_ (Fig. 1*E*).

Next, we aimed to investigate whether Atg18 could cluster liposomes that contain PtdIns(3,4,5)P_3_. Previous studies have shown that Atg18 rapidly forms oligomers when bound to liposomes containing PtdIns(3,5)P_2_ (8, 10), tethering two opposing membranes (9). Using dynamic light scattering (DLS), we showed that Atg18 clusters liposomes containing PtdIns(3)P, PtdIns(3,5)P_2_, and PtdIns(3,4,5)P_3_ (Fig. 1*F*), resulting in larger particle sizes and longer diffusion times for PtdIns(3,5)P_2_- and PtdIns(3,4,5)P_3_-liposomes, indicating increased aggregation.

As previously mentioned, the molecular mechanism underlying the Atg18-polyphosphoinositide binding remains unclear, with conflicting findings regarding its stereoselectivity and binding affinity. Although most studies agree that Atg18 specifically binds to both PtdIns(3)P and PtdIns(3,5)P_2_, there is no consensus on its relative affinities. Some studies suggest a higher affinity for PtdIns(3)P (3, 6, 13), aligning with its role in autophagosome formation, while others identify PtdIns(3,5)P_2_ as the main binding PtdInsP_x_ (4, 9, 11), highlighting its role in endolysosomal membrane regulation and vesicle trafficking. These discrepancies may result from variations in methodologies, such as lipid-binding assays or *in vivo* studies. In this study, we combined a series of well-established *in silico* and *in vitro* methods to study the interaction between Atg18 and PtdInsP_x_. Our findings unambiguously demonstrate that Atg18 binds to PtdIns(3,5)P_2_ with significantly higher affinity than PtdIns(3)P, resolving previous inconsistencies in the literature (3, 4, 6, 9, 11, 13) and in agreement with earlier reports suggesting that PtdIns(3)P alone does not recruit Atg18 to the pre-autophagosomal structure (PAS) (11). Additionally, we provide the first evidence that Atg18 binds to PtdIns(3,4,5)P_3_-containing membranes, and clusters vesicles in a manner similar to PtdIns(3,5)P_2_. This result is supported by previous studies identifying WIPI1, a mammalian homolog of Atg18, as a component of the plasma membrane (14), which is highly enriched in PtdIns(3,4,5)P_3_ (15). Moreover, Atg18 promotes protein transport from early endosomes toward the plasma membrane (16), which has also been suggested to contribute, at least in part, to the formation of the autophagosomal membrane (17). Therefore, our findings raise new questions regarding the functions of PROPPIN and WIPI proteins in regulating intracellular vesicle trafficking and organelle membrane organization, thereby opening new avenues for further research.

## Materials and Methods

For this study, we combined molecular dynamic simulations and biophysical methods, such as isothermal titration calorimetry (ITC), cosedimentation, fluorescence resonance energy transfer (FRET), and dynamic light scattering (DLS). Details of all methodologies applied in this study are included in *SI Appendix*.

## Data, Materials, and Software Availability

Manuscript data have been deposited in Zenodo (https://zenodo.org/records/14779518).

## Acknowledgments

A.P.-L. acknowledges funding by a Ramon y Cajal grant (RYC2018-023837-I) and Max Planck Society through the funding of a Max Planck Partner Group on “Regulation of the SNARE zippering by complexins and synaptotagmins” at the University of Granada. A.D.-A. acknowledges a Garantía Juvenil contract from Consejería de Educación, Juventud y Deporte de la Comunidad de Madrid cofunded by Fondo Social Europeo. M.M.-M acknowledges Centro de Supercomputación de Galicia (CESGA) and CSIC for the use of Finis Terrae III and HPC Drago, respectively.

## Supporting Information for

### Supporting Information (SI) Text

#### Materials and Methods

##### Materials

Key resources table.

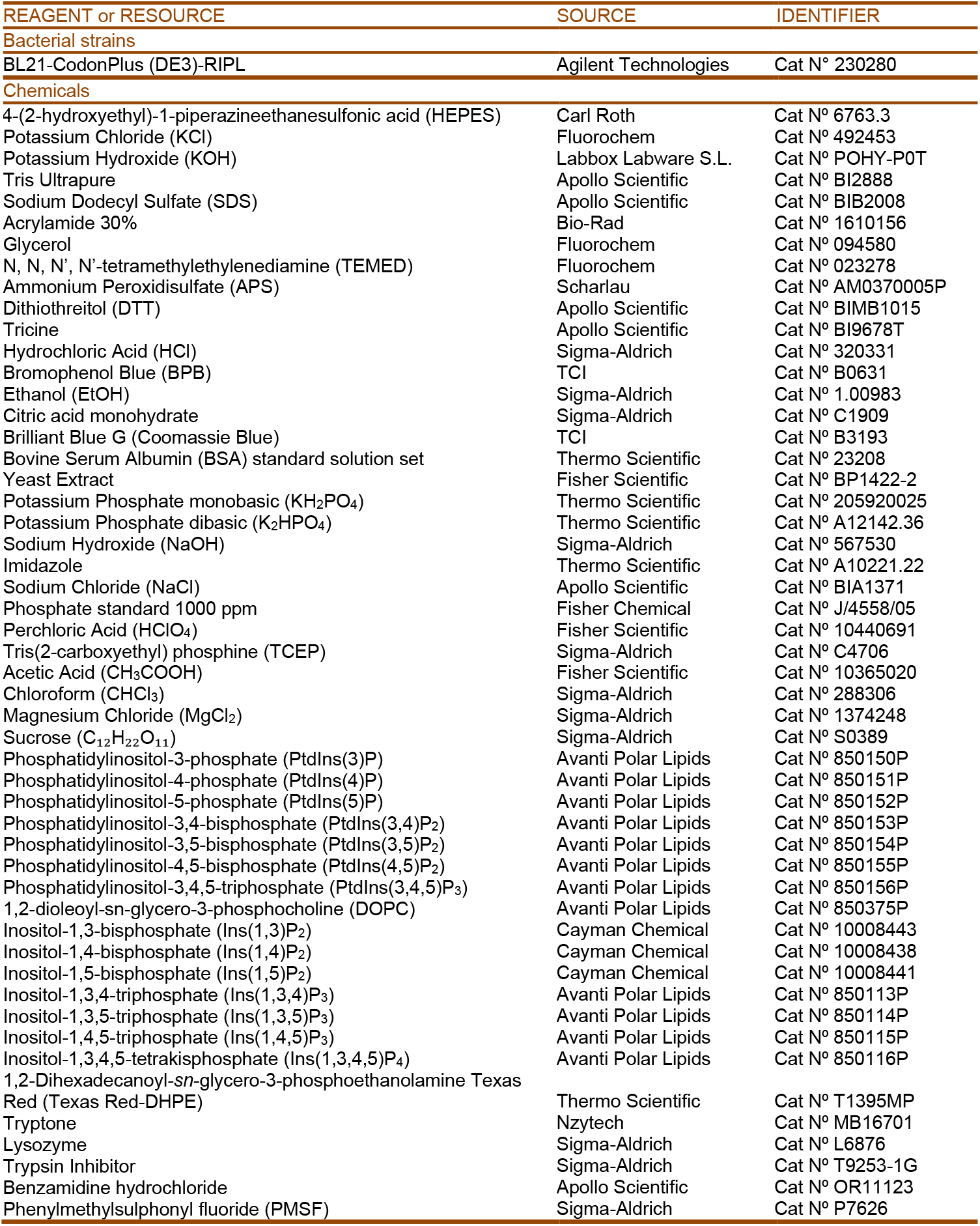
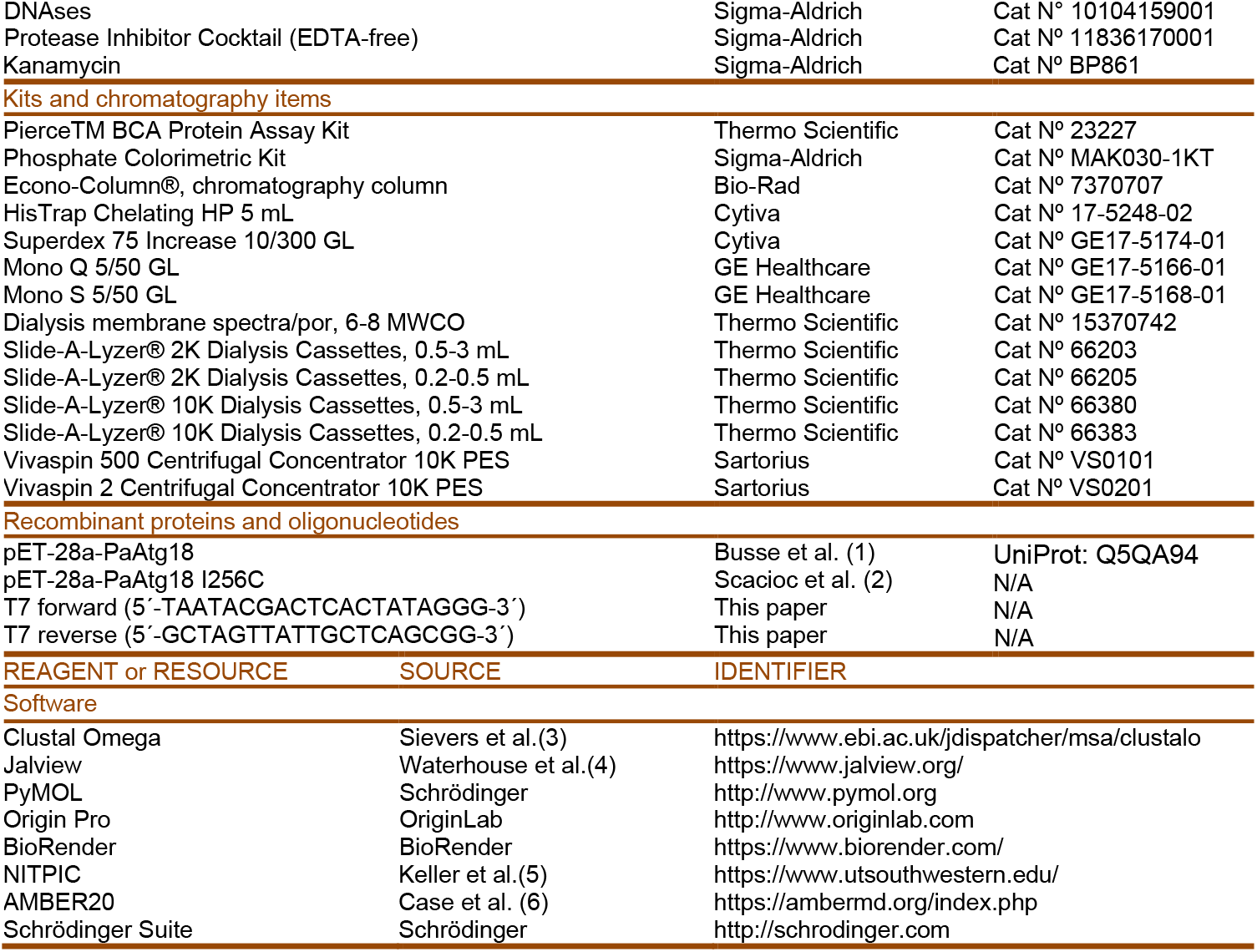

##### Methods

###### Induce Fit Docking Studies

The structure of Atg18 was taken from the PDB structure of code 5LTD (2) corresponding to Atg18 from *Pichia angusta* bound to two phosphate groups. The protein was prepared with the Protein Preparation Wizard tool within the Schrödinger suite of programs (Schrödinger Release 2021-4: Protein Preparation Wizard; Epik, Schrödinger, LLC, New York, NY, 2021; Impact, Schrödinger, LLC, New York, NY; Prime, Schrödinger, LLC, New York, NY, 2021). The ligand, Ins(1,3,4,5)P4, was build using the Maestro interface of Schrodinger suite version 13.0 (release 2021-4), and refined using Schrödinger's LigPrep, with the OPLS4 force field at pH 7.0 ± 2 (Schrödinger Release 2021-4: LigPrep, Schrödinger, LLC, New York, NY, 2021). The Induced fit docking protocol within the Schrödinger suite of programs (Schrödinger Release 2021-4: Induced Fit Docking protocol; Glide, Schrödinger, LLC, New York, NY, 2021; Prime, Schrödinger, LLC, New York, NY, 2021) was used to study the binding of Ins(1,3,4,5)P_4_ to Atg18. Atg18 has two different binding sites for PtdInsP_x_, each contain a histidine residue in the binding site, His221 and His292. Therefore, four IFD simulations were carried out, considering the two binding pockets and the two neutral tautomers of histidine, with the proton on either the delta or epsilon nitrogen. The binding pockets were defined as a cubic box centered on each of the two phosphate molecules bound to Atg18 (PDB code 5LTD). The IFD protocol allows the flexibility of the side chains of the residues around 5 Å from the ligand. First, a soft docking was performed with Glide, with the van der Waals radii (rdW) of the ligand and protein reduced by 50 %. After optimization of the sidechains of the residues within 5 Å from the ligand with Prime, a second docking of the ligands with glide was performed, using the rdW without any reduction. Pymol (PyMol Molecular Graphic System V2.3.3 Schrödinger, LLC) was used for visualization.

###### Molecular dynamic (MD) studies

MD studies were performed with selected IFD poses of Ins(1,3,4,5)P_4_ bound to Atg18, using AMBER 20 and the FF19SB force field.2 Each complex was surrounded by an octahedral box of TIP3P water molecules of 10 Å around the protein in each direction. Periodic boundary conditions were used. Initially, a steepest descent minimization for 30.000 steps was carried out, with a positional restraint weight of 5.0 Kcal mol-1 Å-2. Restraints were gradually weakened and removed in the last 10.000 steps. Next, the system was equilibrated over 125 ps, with an integration time step of 1.0 fs. Initially, water was equilibrated, while the solutes where restrained, with positional restraint weight of 5.0 Kcal mol-1 A-2. Retrains were progressively weakened and removed in the last 25 ps. Temperature was maintained at 300 K using Langevin dynamics. Finally, a production simulation of 500 ns was run, under the NPT ensemble. For this stage the SHAKE algorithm was used and an integration time step of 2.0 fs. The particle mesh Ewald method was used to calculate electrostatic interaction. Three independent replicas were done for each of the complexes. For visualization and creating figures PyMOL (PyMol Molecular Graphic System V2.3.3 Schrödinger, LLC) was used.

###### Protein expression and purification

A synthetic gene encoding *Pichia angusta* Atg18 (*Pa*Atg18, UniProt entry Q5QA94) cloned in the pET-28a vector was used for expression in *Escherichia col*i strain BL21-CodonPlus (DE3)-RIPL as described previously (1). Bacterial pellet of *Pa*Atg18 was resuspended in lysis buffer (30 mM HEPES, 300 mM NaCl, 10 mM MgCl_2_; pH∼7.5) with added protease inhibitor benzamidine, phenylmethylsulfonyl fluoride (PMSF), trypsin inhibitor, and a tablet of cOmplete™, Mini, EDTA-free protease inhibitor cocktail (Roche). A tip of DNAse and lysozyme was also added. The cells were lysed by 3 cycles of 1 minute each (25% amplitude and duty 50), using a digital sonicator S450D (Brandson). Then, the lysate was centrifuge for 20 minutes at 13000 rpm at 4ºC (Type JA-30.50 Ti rotor in Beckman Avanti J-30I Ultracentrifuge) and the supernatant was afterward filtered using a 0.45 μm pore size filter (company). Next, N-terminal His-tagged wild-type *Pa*Atg18 was purified with a 5 mL HisTrap Chelating HP 5 mL column (GE Healthcare) equilibrated with buffer A (30 mM HEPES, 500 mM NaCl, pH= 7.50) and eluted with a gradient from 0 to 100% buffer B (30 mM HEPES, 500 mM NaCl, 1000 mM Imidazole, pH= 7.50) on a ÄKTA system (GE Healthcare). *Pa*Atg18 fractions were collected and dialyzed overnight at 4 °C in dialysis buffer (30 mM HEPES, 200 mM NaCl, 0.1 mM TCEP, pH= 7.50), and subsequent anion exchange chromatography was performed using a 0 mM-1000 mM NaCl gradient in 30 mM HEPES, 0.1 mM TCEP, pH= 7.50 (Mono Q^™^ 5/50 GL, GE Healthcare), followed by size-exclusion chromatography (Superdex 75 Increase 10/300 GL, GE Healthcare) using 30 mM HEPES, 300 mM NaCl, 0.1 mM TCEP, pH 7.50. Protein concentrations were estimated by using a molar extinction coefficient of 29340 M^-1^cm^-1^ and protein purity was checked by SDS-PAGE. Proper protein folding was assessed using circular dichroism (CD).

###### Liposome preparation for cosedimentation binding assay

Large unilamellar vesicles (LUVs) were composed of 2% of the different physiological phosphatidylinositol (PInsP_x_) and 98% of 1,2-dioleoyl-sn-glycero-3-phosphocholine (DOPC). Lipids mixtures were dried in glass tubes with an oxygen-free nitrogen stream and kept under vacuum overnight in a vacuum desiccator. Lipid films were resuspended in buffer (25 mM HEPES, pH 7.5, 150 mM KCl, and 200 mM sucrose) by vigorous vortexing. LUVs with a diameter of around ∼140 nm were prepared by extrusion of the rehydrated phospholipid suspensions with 21 strokes through 0.1 μm polycarbonate membranes (Millipore, Inc.) with a miniextruder (Avanti Polar Lipids), and then dialyzed against the reaction buffer (25 mM HEPES, pH 7.5, and 150 mM KCl). The homogeneity of liposome preparations was confirmed by dynamic light scattering measurements with a Zetasizer μV equipment (Malvern Panalytical). Total phospholipid concentration was determined using a phosphate colorimetric assay kit (Sigma-Aldrich).

###### Liposome preparation for fluorescence-based liposome binding assay

Lipid mixtures containing 2% of the different physiological phosphatidylinositol (PInsP_x_), 96% of 1,2-dioleoyl-sn-glycero-3-phosphocholine (DOPC), and 2% of Texas Red-PE were prepared as described above. Lipid films were resuspended in buffer (25 mM HEPES, pH 7.5 and 150 mM KCl) by vigorous vortexing and then extruded 21 strokes through 0.1 μm polycarbonate membranes. The homogeneity of liposome preparations and total lipid concentration were determined as explained above.

###### Protein labelling

Alexa Fluor 488 C5 Maleimide (Thermo Fisher Scientific) was used for specific labelling of the cysteine mutant *Pa*Atg18 I256C. *Pa*Atg18 I256C was incubated with a fivefold molar excess of the dye while gently shaking overnight at 4 °C. Unreacted dye was removed from the labeled protein with a HiTrap desalting column (Cytiva) using 25 mM HEPES, pH 7.5, and 150 mM KCl. Protein concentration was monitored using Pierce^™^ BCA protein assay kit (Thermo Scientific). The labelling efficiency of Atg18 was higher than 90%.

###### Cosedimentation binding assay

1 μM of *Pa*Atg18 was incubated with LUVs containing sucrose (300 μM total lipid in 100 μl) in buffer containing 25 mM HEPES, pH 7.5, and 150 mM KCl at 25°C for 30 min. Membrane-bound *Pa*Atg18 was separated from free *Pa*Atg18 by centrifuging the mixture at 13,000 rpm for 80 min at 20 °C (Type 12094 Sigma rotor in Sigma 1-14K centrifuge). Aliquot *Pa*Atg18 detection and quantification were performed by densitometry analysis of Coomassie blue stained SDS-PAGE gels using a ChemiDoc Touch scanner (BIORAD). Two biological repeats with two technical repeats each were done for all conditions.

The analysis of SDS gels was carried out using Fiji (7) with a custom macro developed in-house. Briefly, the process starts with the subtraction of the background signal. To achieve this, a new background image is generated by applying a strong median filter. This filter effectively removes positive regions while preserving the background, allowing for the isolation of the gel signal. The resulting background-free image, referred to as the “Gel image”, is obtained by subtracting the filtered background from the original raw image. Once the background has been removed, an automatic threshold is applied using the Triangle algorithm in Fiji. This step generates a binary image, known as the “ROI image”, which highlights the positive regions of the gel with a value of 1, while the rest of the image is set to 0. The binary ROI image is then used to extract data from the Gel image. The Gel image retains background-subtracted values in the positive regions, ensuring that only relevant areas are analyzed. These selected regions are subsequently processed using the “Analyze Particles” tool in Fiji, which allows for the measurement of the average pixel intensity and the size of each positive region. This approach provides a consistent and reproducible method for quantifying the bands observed in SDS gels, ensuring accurate analysis of the experimental data. Standard deviations (SD) were calculated.

macro “Gels []” {

setTool(“rectangle”);

waitForUser(“Waiting”, “Manually select the rectangle and click OK”);

run(“Duplicate…”, “title=Gel”);

run(“32-bit”);

// If the bands are not illuminated, use the following command:

run(“Reciprocal”);

// Creating the background Image (named Median)

run(“Duplicate…”, “title=ROI”);

run(“Duplicate…”, “title=Median”);

run(“Median…”, “radius=50”);

// Subtracting the background

imageCalculator(“Subtract”, “ROI”,”Median”);

imageCalculator(“Subtract”, “Gel”,”Median”);

// Preparing the image for the ROI selection. selectImage(“ROI”);

run(“Square Root”);

run(“Square”);

run(“Remove Outliers…”, “radius=5 threshold=20 which=Bright”);

run(“Gaussian Blur…”, “sigma=3”);

// Automated ROI selection.

setAutoThreshold(“Triangle dark”);

setOption(“BlackBackground”, false)

run(“Convert to Mask”);

//run(“Divide…”, “value=255.0000”);

imageCalculator(“Multiply”, “Gel”,”ROI”);

setThreshold(0.1, 1000000000000000000000000000000.0000);

run(“NaN Background”);

// Analyzing particles.

electImage(“Gel”);

run(“Analyze Particles…”, “size=300-Infinity pixel exclude clear overlay add”);

}

###### Fluorescence-based binding assay

LUVs for fluorescence-based binding assay were prepared as described above. Alexa Fluor 488 fluorescence emission was measured at 25 °C with a Jasco FP-8500 spectrophotometer (JASCO UK Limited) in the range of 510 nm to 700 nm with an excitation wavelength of 480 nm and 2.5 nm slit width using 1 μM protein (Alexa Fluor 488-*Pa*Atg18 I256C) and 300 μM total lipid of different lipid mixtures. Total phospholipid concentration was determined using a phosphate colorimetric assay kit (Sigma-Aldrich).

Fluorescence resonance energy transfer (FRET) between Alexa Fluor 488-labelled Atg18 and Texas Red-DHPE was monitored by the decrease of the Alexa Fluor 488 fluorescence emission at 519 nm and analyzed using the equation:

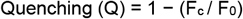

where *F*_*c*_ and *F*_0_ are the fluorescence emission at 519 nm in the presence and in the absence of liposomes containing the FRET acceptor Texas Red-DHPE, respectively. Quenching at different liposome concentrations was analyzed by a Hill equation:

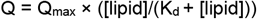

where Q_max_ represents the calculated maximal quenching, [lipid] the lipid concentration and K_d_ the equilibrium dissociation constant, using Origin 2020 software (Origin Labs Inc., Northampton, MA, USA). Three technical repeats were done for all proteins both in the presence and in the absence of vesicles. Standard deviations (SD) were calculated.

###### Aggregation assay

*Pa*Atg18-induced liposome aggregation was monitored by dynamic light scattering measurements with a Zetasizer μV equipment (Malvern Panalytical). Liposome compositions were similar to those of the liposomes used for fluorescence-based binding assay. Thus, 1 μM of *Pa*Atg18 was mixed with liposomes (300 μM total) and incubated in buffer containing 25 mM HEPES, pH 7.5, and 150 mM KCl (100 μl final volume) at 25 °C for 30 min. Two biological repeats with two technical repeats each were carried out for all conditions. Standard deviations (SD) were calculated.

###### Isothermal titration calorimetry (ITC)

ITC experiments were carried out as previously described (8). Titration of ∼25 μM of *Pa*Atg18 with different physiological phosphatidylinositols (PtdInsP_x_) head groups (InsP_x_) were performed at 25 °C in 50 mM HEPES, pH 7.5, and 150 mM KCl. To obtain the effective heat of binding, results of the titration were corrected using buffer-protein and ligand-buffer controls. Finally, NITPIC (5) and Origin (Origin Labs Inc.) software packages were used to analyze these data. Two biological replicates, each with two or three technical replicates, were conducted for all conditions. Standard deviations (SD) were calculated.

###### CD spectroscopy

CD measurements were performed using a J-715 spectropolarimeter (JASCO UK Limited) for ∼0.2 mg/ml of protein (monitored the absorbance at 280 nm using a NanoDrop 2000, Thermo Scientific) in 0.15 M NaF, 20 mM NaH_2_PO_4_ pH 7.5 —in a 0.1 cm Hellma quartz cuvette— between 200 and 260 nm at 25 °C, using 1 nm step size and 2 nm band width. Temperature scan was performed at 216 nm from 40 °C to 70 °C with a heating rate of 1 °C/min with a temperature step of 0.2 °C. Bandwidth was 2 nm and the averaging time was 16 s. Samples were centrifuged at 14000 rpm before measurements to remove aggregated and precipitated protein.

###### Declaration of generative AI and AI-assisted technologies in the writing process

During the preparation of this work, the authors used OpenAI’s ChatGPT to refine the language and grammar of this manuscript. The authors reviewed and confirmed all AI-generated content for accuracy and compliance with ethical standards. The authors take full responsibility for the content of the publication.

## Notes

### Competing Interest Statement

The authors have declared no competing interest.

### Summary of Updates

Change the paragraph from: 'Furthermore, Atg18 promotes protein transport from early endosomes toward the plasma membrane (16), and the autophagosomal membrane has been suggested to derive, at least in part, from the plasma membrane (17).' To: 'Moreover, Atg18 promotes protein transport from early endosomes toward the plasma membrane (16), which has also been suggested to contribute, at least in part, to the formation of the autophagosomal membrane (17).'" Additionally, remove Table 1 to comply with PNAS requirements.

